# Quantitative Analysis of External Urethral Sphincter Stimulation Parameters for Modulating Urinary Output

**DOI:** 10.64898/2026.05.08.723239

**Authors:** Yifan Wang, Md Abdul Kader Tushar, Orion Lucero, Philippe E. Zimmern, Zhengwei Li

**Affiliations:** Department of Biomedical Engineering, University of Houston, USA; Department of Urology, University of Texas Southwestern, USA; Department of Biomedical Engineering and Department of Biomedical Sciences at University of Houston, USA

**Keywords:** Neuromodulation, Electrical stimulation parameters, Neurogenic lower urinary tract dysfunction, external urethral sphincter, Rehabilitation engineering.

## Abstract

**Objective:** Neurogenic lower urinary tract dysfunction (NLUTD) impairs bladder control and remains difficult to treat. We aim to define how electrical stimulation (ES) parameters of the external urethral sphincter (EUS) affect urinary leakage thresholds to guide neuromodulation strategies for NLUTD.

**Methods:** We performed direct EUS stimulation in anesthetized rats using charge-balanced biphasic pulses while systematically varying current amplitude (0.5–3.0 mA), frequency (20–100 Hz), and pulse duration (0.5–3 ms). Urine leakage thresholds were mapped across the multidimensional parameter space.

**Results:** Stimulation parameters exhibited strong nonlinear interdependence in determining leakage onset. At a fixed pulse duration, higher current amplitudes required lower stimulation frequencies to evoke leakage. Increasing pulse duration substantially reduced both current and frequency thresholds. Age and sex caused modest shifts in absolute thresholds but did not alter the fundamental parameter–response relationships.

**Conclusion:** Pulse duration, current amplitude, and frequency jointly govern urinary leakage thresholds, with pulse duration serving as the dominant modulator of stimulation efficiency.

**Significance:** This work establishes a quantitative framework for charge-efficient stimulation parameter selection, enabling the design of energy-aware, precision neuromodulation protocols and implantable systems for NLUTD rehabilitation.

## I. INTRODUCTION

NEUROGENIC lower urinary tract dysfunction (NLUTD) arises from common neurological disorders such as spinal cord injury (SCI), Parkinson’s disease, multiple sclerosis and other neurodegenerative diseases. NLUTD impairs bladder function, and the inability to empty the bladder can lead to serious complications such as urinary tract infections (UTIs), hematuria, bladder stones, and even bladder cancer [1]. Current management strategies primarily include intermittent catheterization, pharmacological treatments, surgical interventions like sphincterotomy and urethral stents [2,3]. However, these methods often have limited efficacy, significant side effects, and poor long-term outcomes, which highlight the need for improved therapeutic solutions.

Neuromodulation strategies targeting the lower urinary tract have been explored for treating urinary disorders. Historically, stimulation efforts focused on the larger or more accessible neural targets, such as pelvic nerve [4], pudendal nerve [5], sacral nerve [6], and even the spinal cord [7]. However, modulating the activity of these nerve targets is challenging since it is difficult to identify the reliable branching of the terminal axons in the densely innervated bladder [8]. Moreover, broad nerve stimulation is non-specific and may indiscriminately block or activate multiple pathways, potentially leading to undesirable side effects. Some recent studies have shifted focus to smaller, more distal nerve pathways (e.g., postganglionic bladder branches) that might offer more specific control, but these tiny nerves are difficult to visualize and manipulate surgically, therefore increasing the risk of inadvertent injury and bladder dysfunction [9].

Accordingly, direct bladder stimulation using implantable electrodes has been proposed as a treatment of NLUTD [10-12]. However, the bladder is a highly dynamic organ that undergoes substantial changes in volume and shape during the filling and voiding, which needs soft and stretchable scaffolds to implant [13]. In addition, most existing studies primarily focus on evoking detrusor contraction but do not adequately address the role of the EUS. The EUS is a critical component of the lower urinary tract, providing active urethral closure during bladder filling to maintain continence [14]. Recent studies focused on using electrical neuromodulation to target the EUS to treat NLUTD. However, the physiological consequences of direct EUS stimulation have been inconsistent in prior studies.

While some studies report that EUS activation via electrical stimulation enhances outlet resistance and improves continence [15], others have found that similar stimulation can paradoxically trigger bladder contraction or urine leakage [16, 17]. These conflicting outcomes suggest that stimulation parameters, such as current amplitude, frequency, and pulse duration will determine whether EUS stimulation reinforces continence or induces a voiding reflex.

The underlying mechanism involves complex neural circuits spanning the spinal cord and brainstem. During normal urine storage, the pontine storage center exerts inhibitory control over parasympathetic outflow from the sacral spinal cord, while activating somatic efferents via the pudendal nerve to maintain EUS contraction [18]. Electrical stimulation of the EUS activates pudendal afferents, which project to spinal interneurons that modulate parasympathetic neurons. Under certain conditions, this modulation may reduce inhibitory tone and facilitate detrusor activation, leading to partial or full micturition. Thus, direct EUS stimulation can either augment continence or invoke a bladder voiding reflex, depending on stimulation parameters and neural state.

Despite these insights, there has been a lack of systematic characterization of the stimulation thresholds and parameter combinations that produce urine leakage (or other bladder responses) via EUS stimulation. To address this gap, the present study uses flexible, implantable electrodes stimulating EUS and systematically investigated how stimulation amplitude, frequency, and pulse duration interact to induce urine leakage in an anesthetized rat model (Fig. 1(a)). We also studied the influence of age and sex on these relationships by mapping the EUS stimulation parameter space and identifying leakage threshold boundaries. We aimed to establish a predictive framework for safe and effective neuromodulation. These findings will inform individualized stimulation strategies and guide the design of implantable systems for managing chronic urinary disorders.

**Fig. 1.**
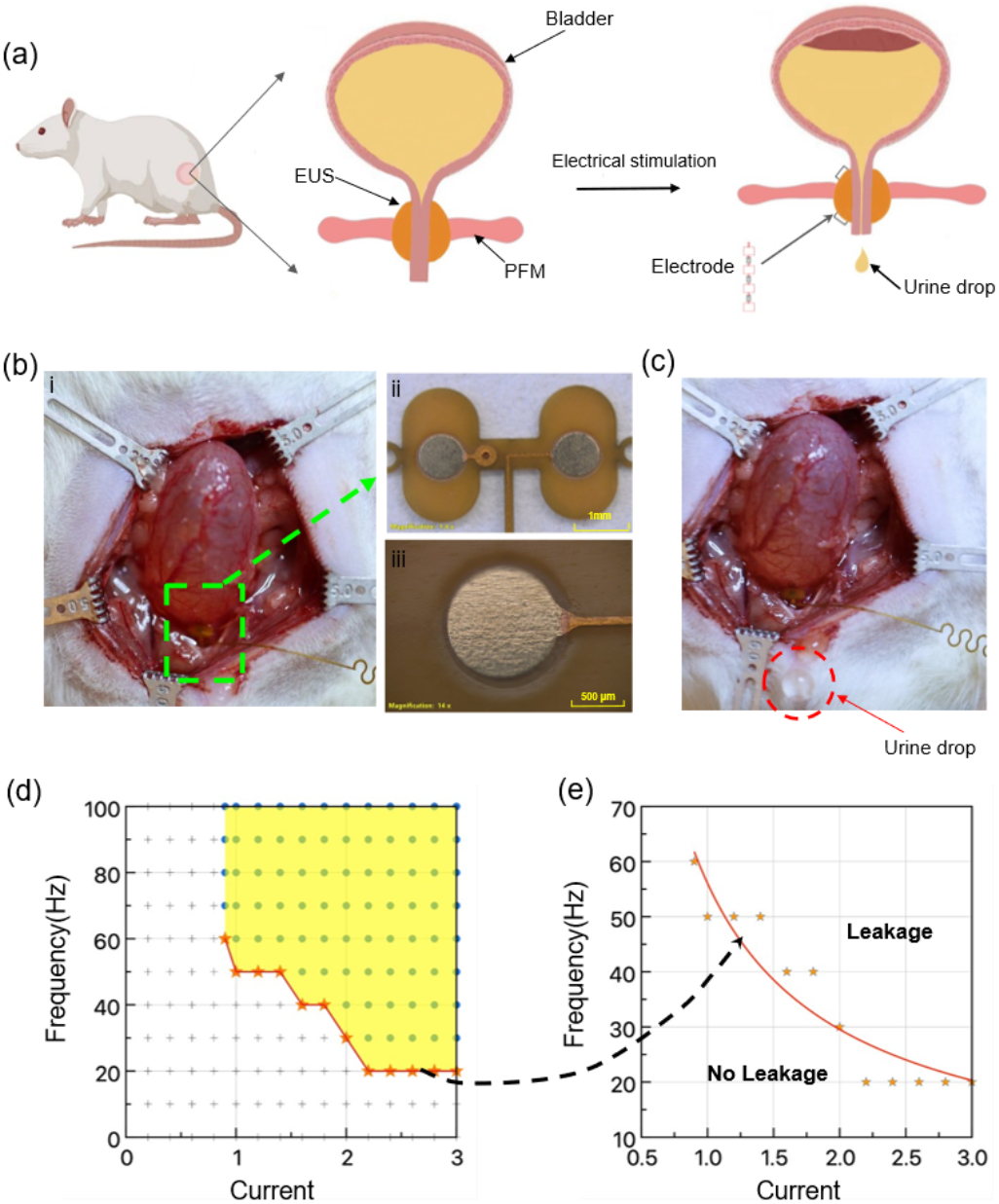
Experimental setup for EUS stimulation and threshold mapping. (a) Schematic illustration of direct electrical stimulation of the EUS in rats to assess urinary outcomes. (b) photograph showing the flexible PCB electrode implanted on the rat EUS. (c) Experimental image showing urine leakage during EUS electrical stimulation. (d) Example map of stimulation outcomes across frequency (20–100 Hz) and current amplitude (0–3 mA) at a fixed 1 ms pulse width. Yellow shading indicates frequency–current combinations that evoked urine leakage. (e) Enlarged view of the threshold boundary from (d), showing the minimum frequency required to induce leakage at each current level (orange star markers). The red curve is a fitted threshold line separating no-leakage versus leakage conditions.

## II. METHOD

### A. Animal Preparation and Surgical Procedure

Adult Sprague–Dawley rats (weight 280–500 g; n = 7; 4 females, 3 males) were obtained from Taconic Biosciences (Germantown, NY). The animals were maintained at controlled temperatures with humidity of 40-65% and had ad libitum access to a standard laboratory diet and water. The animals were anesthetized using isoflurane (2%) in a constant oxygen flux (0.5 L/min). Body temperature was maintained at approximately 37°C using a warming pad, and heart rate and respiration were continuously monitored. All experimental procedures were reviewed and approved by the University of Houston Institutional Animal Care and Use Committee (IACUC) under Protocol No. PROTO202100048. All animal work was performed in strict accordance with the committee’s ethical guidelines and federal regulations.

All animals were placed in a supine position with limbs gently secured to the operating table. The ventral abdomen was shaved, followed by three sequential scrubs of povidone iodine.

A low midline laparotomy of 2-2.5 cm was performed to expose the bladder (Fig. 1(b)(i)). A PE-50 catheter was inserted through a small incision at the bladder dome and secured with a 6-0 silk suture to prevent leakage. The catheter was connected to a syringe pump for repeated process. The urine leakage was video recorded and analyzed off-line.

### B. Flexible Stimulation Electrodes

The flexible printed circuit board (PCB) electrode was designed using Fusion 360 and fabricated in panelized form by an external manufacturer (PCBWay). The electrode was realized on a flexible polyimide–copper laminate consisting of a 75 µm thick of polyimide (PI) substrate and an 18 µm thick of copper (Cu) layer (Fig. 1(b)(ii) and (iii)). Optical microscopy images at low and high magnification confirmed the well-defined circular electrode area and the integrity of the copper interconnect without visible cracks or delamination. The PCB electrode comprised circular copper stimulation pads interconnected by serpentine copper traces to enhance mechanical flexibility and strain tolerance (Supplementary Fig. 1(a)). Each serpentine interconnect terminated at a dedicated connection pad located at the edge of the PCB. Electrical interfacing was achieved by soldering insulated copper wires to the connection pads, providing a robust and low-resistance contact (Supplementary Fig. 1(b)). The soldered copper wires were subsequently connected to the output terminals of the pulse stimulator using hook electrodes, enabling reliable bipolar stimulation during experiments (Supplementary Fig.2). The serpentine copper interconnects exhibited excellent stretchability, accommodating up to 40% elongation (from 68 mm to 95.2 mm) without connection damage, as demonstrated in the before- and after-stretching images (Supplementary Fig. 3).

### C. Stimulation Setup

Each flexible PCB electrode was placed directly on the external urethral sphincter (EUS) using a high-magnification surgical microscope to ensure accurate positioning. Electrode placement was guided by anatomical landmarks in the pelvic floor musculature. The electrode was connected to an isolated constant-current bipolar stimulator (Model 2100, A-M Systems, Sequim, WA). All stimulations were delivered using biphasic, charge-balanced rectangular pulses to avoid net direct current accumulation at the electrode–tissue interface. The stimulator supported full programmability of pulse amplitude, duration, and frequency. Stimulation parameters included pulse durations of 0.5 ms, 1 ms, and 3 ms; current amplitudes ranging from 0.5 to 3.0 mA; and pulse repetition frequencies between 20 and 100 Hz. Both single-pulse and repetitive pulse-train protocols were used. Rest periods were

### D. Parameter Mapping and Outcome Assessment

To characterize the interaction between stimulation parameters and urinary responses, we systematically mapped leakage thresholds by varying current amplitude from 0.5 mA to 3.0 mA (in 0.5 mA increments) and frequency from 20 to 100 Hz (in 10Hz increments) at fixed pulse durations ranging from 0.5 to 3.0 ms (in 0.5 ms increments). For each parameter combination, stimulation was applied for 2–5 seconds, and the presence or absence of visible urine leakage from the urethral meatus was recorded (Fig. 1(c)). All mapping experiments were repeated at least twice per condition to confirm consistency. Results were organized into 2D parameter maps, with shaded regions indicating frequency–current pairs that triggered leakage (Fig. 1(d)). To quantify the threshold boundary, we extracted the minimum frequency required to induce leakage at each tested current and plotted threshold curves for comparison across conditions (Fig. 1(e)). These experiments were performed in all seven rats to assess inter-animal variability and reproducibility. Age- and sex-related influences were also evaluated by comparing threshold curves across three female rats (ages 9, 10, and 12 months) and three male rats (ages 4, 6, and 8 months). All tests were performed under consistent experimental conditions, and trends were analyzed to determine whether age or sex affected the shape or position of the leakage threshold curves.

## III. RESULTS

### A. Mapping Stimulation Parameters to Urinary Outcomes

We systematically mapped the stimulation parameter space to examine how current amplitude, frequency, and pulse duration interact to elicit urine leakage during EUS stimulation. In a representative rat, we examined the effect of pulse duration on the frequency–current threshold for leakage. Using a short pulse width of 0.5 ms, we mapped the combinations of current (0.5–3.0 mA) and frequency (20–100 Hz) that elicited urine leakage (Fig. 2(b)). At 0.5 ms, the “leakage window” was relatively narrow: leakage only occurred at the higher end of the stimulation parameters. Specifically, a minimum of approximately 1.5 mA and ≥60 Hz was required to initiate leakage, and even at the maximum current of 3 mA, frequencies above ∼40 Hz were still needed. These observations indicate that very brief pulses did not deliver enough charge per pulse at low currents, necessitating both high amplitude and high frequency to overcome the threshold for inducing a leak.

**Fig. 2.**
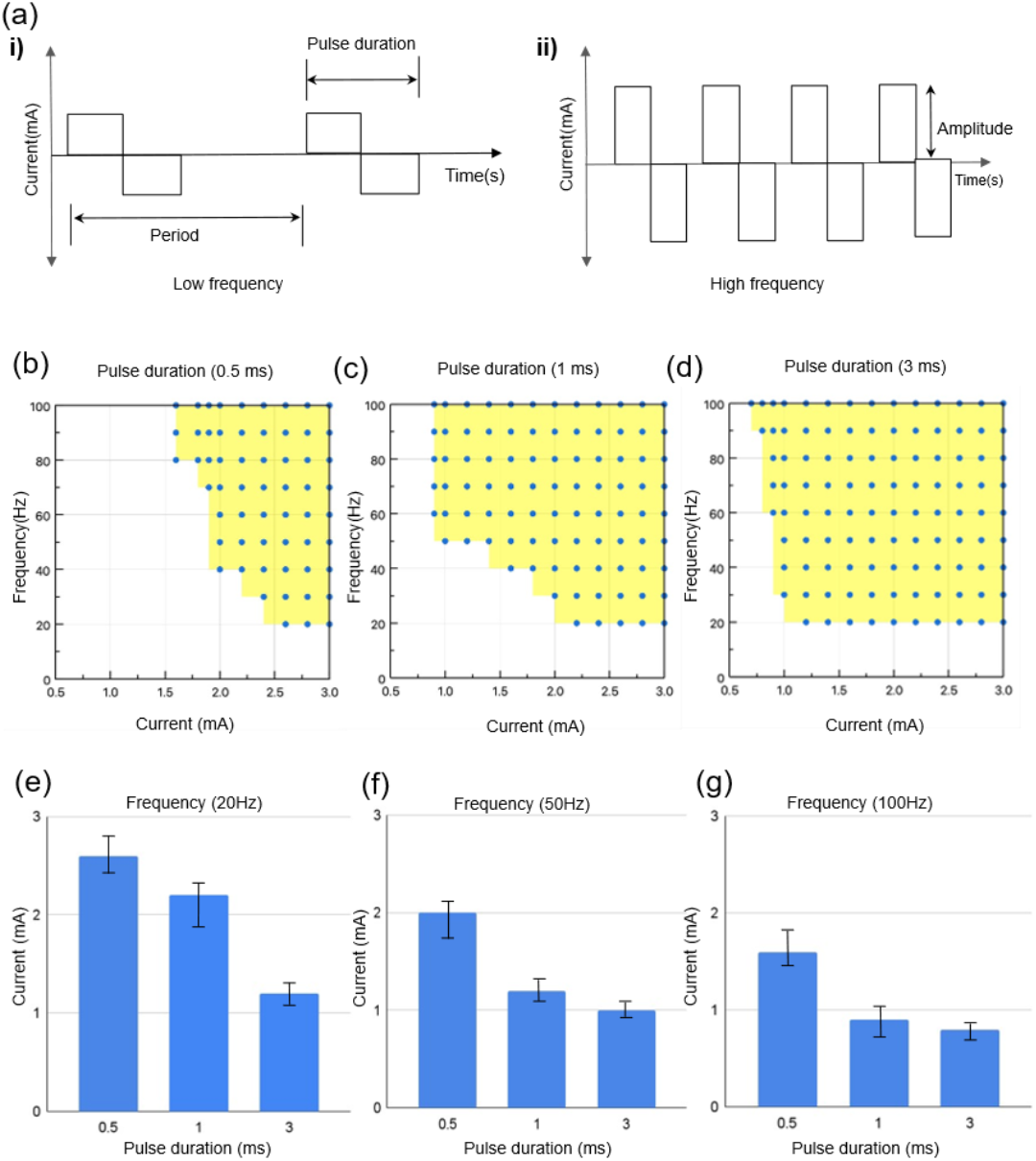
Effects of EUS stimulation parameters on the urinary outcomes. (a) Representative biphasic stimulation waveforms with indicated different parameters, at (i) a low and (ii) high frequency (short interpulse interval). (b-d) Frequency–current response maps for pulse durations of 0.5 ms, 1 ms, and 3 ms respectively. Yellow regions denote combinations of stimulation frequency and current that produced urine leakage (‘leakage window’). (e–g) Current thresholds for leakage as a function of pulse duration at fixed stimulation frequencies of 20 Hz, 50 Hz, and 100 Hz respectively.

We next repeated the test using 1 ms pulses (Fig 2(c)) and noted that the effective parameter space for leakage expanded substantially. Leakage occurred at lower currents (1.0–1.2 mA) when paired with moderate-to-high frequencies (≥60 Hz), while at higher currents (e.g., 2.0–2.5 mA), the minimum frequency required for leakage dropped to 20–30 Hz. The enlarged yellow region and downward shift in the boundary confirmed a synergistic interaction between current amplitude and pulse duration, where increasing pulse width reduces the frequency needed to reach threshold. By using 3 ms pulses (Fig 2 (d)), leakage was observed across nearly all tested combinations except at the lowest amplitudes and frequencies. Even at 0.8–1.0 mA, leakage could be elicited at moderate frequencies (40–50 Hz), and above 1.5 mA, leakage occurred across most of the frequency ranges. The expanded yellow region indicates that longer pulses markedly improve stimulation efficiency, lowering both the amplitude and frequency requirements to initiate leakage.

We further investigated the relationship between pulse duration and current threshold for urine leakage under fixed stimulation frequencies (20 Hz, 50 Hz, and 100 Hz). At 20 Hz (Fig.2 (e)), the shortest pulse (0.5 ms) demanded the highest current (2.6 mA), while the longest pulse (3 ms) reduced the threshold to nearly half (1.2 mA). At 50 Hz (Fig. 2(f)), the same trend persisted, with thresholds dropping from 2.0 mA (0.5 ms) to 1.0 mA (3 ms). At 100 Hz (Fig. 2(g)), overall current requirements were lower, but the pattern remained consistent: 1.6 mA for 0.5 ms, 0.9 mA for 1 ms, and 0.7 mA for 3 ms. These findings confirmed that longer pulse durations significantly reduce amplitude demands, even at low frequencies. This indicates that pulse duration is a critical parameter for optimizing stimulation protocols, enabling lower currents and potentially reducing tissue stress while maintaining functionality.

### B. Effect of Gender and Age

To assess the influence of biological variability, we mapped leakage thresholds in multiple rats of different ages and sexes. Fig. 3(a)–(c) show results for three female rats aged 9 months, 10 months, and 12 months, respectively. All three females exhibited the same qualitative pattern: longer pulse durations shifted the leakage threshold curves downward (requiring lower frequencies at a given current) compared to shorter pulses. In the 9-month-old female (Fig. 3(a)), for example, a 0.5 ms pulse required frequencies around 100 Hz at low currents (∼1.5 mA) and still about 50 Hz at 3 mA to cause leakage. By contrast, with a 3 ms pulse, leakage occurred at much lower frequencies (around 50 Hz at 1 mA, and ∼20 Hz at 2.5 mA). The 10-month female (Fig. 3(b)) showed a nearly identical profile to the 9-month rat, with only minor differences in the exact threshold points. The 12-month female (Fig. 3(c)) had slightly higher thresholds overall: even at the maximum current of 3 mA, the 0.5 ms pulses still required ∼60 Hz to induce leakage, and the curves for 1 ms and 3 ms pulses were shifted a bit upward compared to the younger rats.

**Fig. 3.**
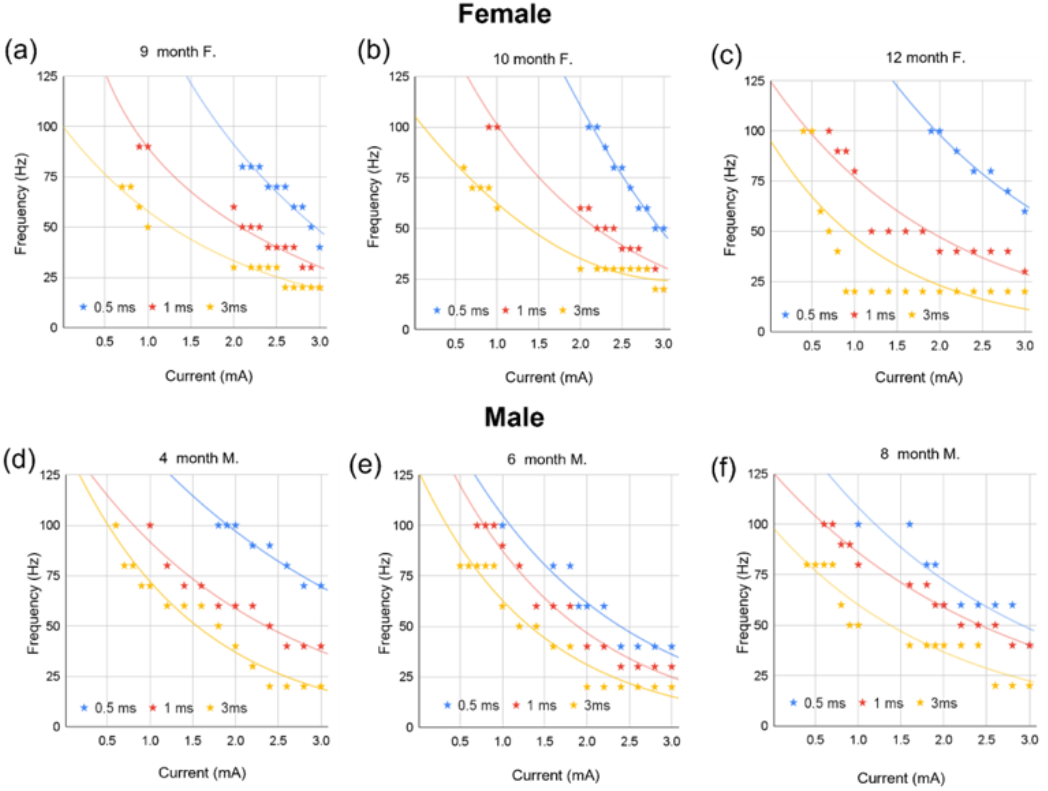
Leakage threshold curves for different ages and sexes. (a–c) Frequency–current leakage threshold curves in female rats aged 9 months (a), 10 months (b), and 12 months (c). Each panel includes curves for three pulse durations (0.5 ms, 1 ms, 3 ms). (d-f) Leakage threshold curves in male rats aged 4 months(d), 6 months(e), and 8 months(f).

A similar analysis in male rats (ages 4 months, 6 months, and 8 months) yielded comparable results (Fig. 3(d)–(f)). In each male, the 0.5 ms pulse condition demanded the highest frequencies for leakage (often near 100 Hz at lower currents), whereas the 3 ms pulses allowed leakage at much lower frequencies. The 4-month-old male (Fig. 3(d)) had a threshold curve comparable to those of the young females: 0.5 ms pulses needed ∼100 Hz at 1 mA (decreasing to ∼70 Hz at 3 mA), while 3 ms pulses brought those requirements down to ∼50 Hz at 1 mA and ∼20 Hz at 2.5 mA. The 6-month male (Fig. 3(e)) showed a slight downward shift in the curves, suggesting a minor reduction in thresholds with maturation (for instance, at 0.5 ms, ∼100 Hz at 1 mA and ∼40 Hz at 3 mA were sufficient to cause leakage). The oldest male (8 months, Fig. 3(f)) had threshold curves that were marginally higher again (e.g., ∼100 Hz was still required at 1 mA for 0.5 ms pulses). Despite these small variations, all male rats demonstrated the same key trend: increasing pulse duration dramatically lowered the frequency (and current) needed to provoke leakage. No fundamental differences were observed between sexes or across age groups; both factors introduced only minor shifts in leakage thresholds without altering the overall parameter–response relationships.

### C. Current–Frequency Threshold Across Animals

To further generalize the findings, we compared stimulation thresholds across all seven rats at a pulse width of 1 ms by isolating the effects of current and frequency. For the current effect, frequency thresholds were measured at fixed currents of 0.5 mA, 1 mA, and 3 mA. At 0.5 mA (Fig 4(a)), all animals failed to elicit leakage with frequency as high as 100Hz, with no meaningful difference between sexes, indicating that low current alone is insufficient to overcome urethral closure pressure. At 1 mA (Fig 4(b)), the urine leakage thresholds decreased substantially, ranging from 80–100 Hz in both females and males, suggesting that moderate current improves efficiency but still demands high-frequency stimulation. At 3 mA (Fig 4(c)), thresholds dropped dramatically to 20–30 Hz in females and 30-40 Hz in males, confirming a strong inverse relationship between current and frequency. Females tended to require slightly lower frequencies than males at high current, indicating marginally greater sensitivity.

**Fig. 4.**
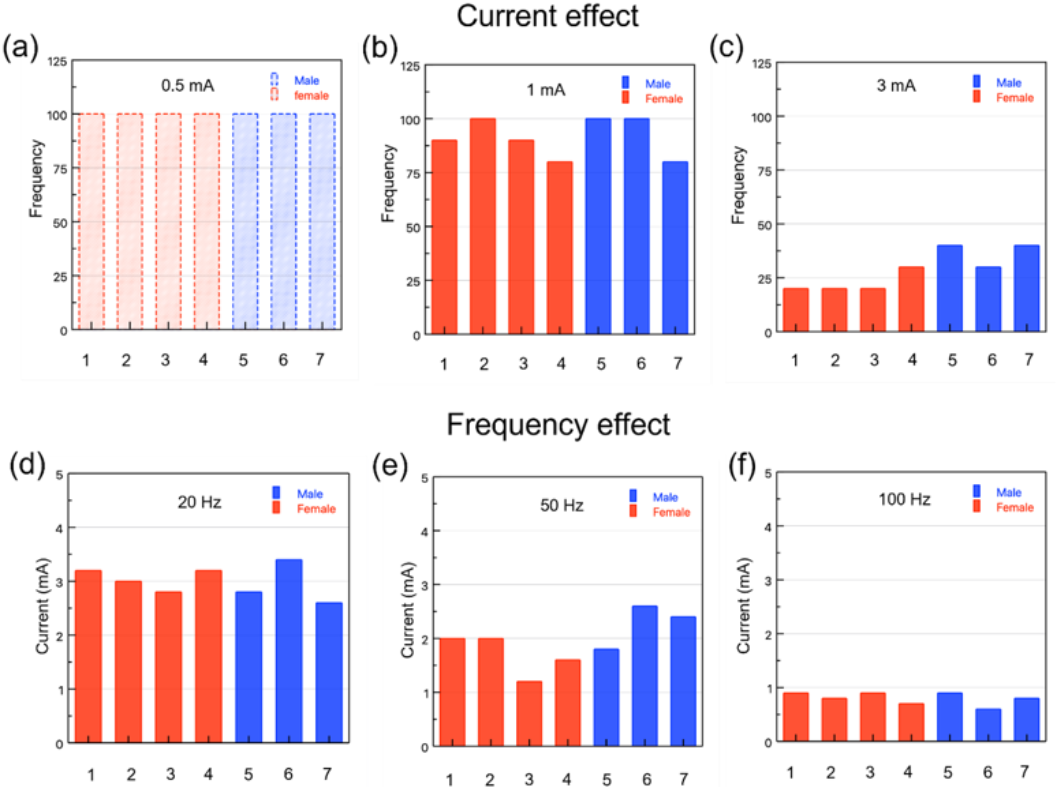
Influence of current and frequency on leakage threshold across animals. (a–c) Minimum stimulation frequency required to induce urine leakage at fixed current amplitudes of 0.5 mA (a), 1.0 mA (b), and 3.0 mA (c). No leakage was observed at 0.5 mA in any animal, even at 100 Hz. At 1.0 mA, leakage required high frequencies (∼80–100 Hz). At 3.0 mA, leakage occurred at substantially lower frequencies (∼20–40 Hz), with female rats generally responding at the lower end of this range. (d–f) Minimum stimulation current required to induce leakage at fixed frequencies of 20 Hz (d), 50 Hz (e), and 100 Hz (f). Current thresholds decreased with increasing frequency, with the lowest thresholds observed at 100 Hz.

We then evaluated the “frequency effect” by holding the stimulation frequency fixed and finding the minimum current that induced leakage at that frequency. For the frequency effect, current thresholds were measured at fixed frequencies of 20 Hz, 50 Hz, and 100 Hz. At 20 Hz (Fig 4(d)), currents were the highest which reflect the difficulty of achieving leakage at low frequency without high amplitude. When frequency increased to 50 Hz (Fig 4(e)), urine leakage thresholds decreased to 2–3 mA, and at 100 Hz (Fig 4(f)), currents were lowest overall (0.6–0.9 mA), indicating that increasing frequency significantly reduces amplitude requirements.

## IV. DISCUSSION

This work systematically characterizes how stimulation amplitude, frequency, and pulse duration interact to determine urine leakage thresholds during direct EUS stimulation. Prior investigations have reported variable functional outcomes following urethral or pudendal stimulation, including both bladder inhibition and bladder activation.

Our findings support the concept that these divergent responses are strongly dependent on stimulation parameter selection and the specific reflex pathways recruited. Woock et al. demonstrated that intraurethral electrical stimulation activates two distinct pudendal afferent pathways that evoke different bladder responses depending on stimulation frequency and anatomical location [19]. Specifically, stimulation of afferents in the dorsal penile nerve (DNP) and cranial sensory nerve (CSN) can engage either spinal or supraspinal reflex circuits, producing excitatory or inhibitory bladder effects. Importantly, bladder responses in that study were frequency-dependent and pathway-specific. Our findings are consistent with this framework in that altering stimulation parameters changes the functional output, suggesting differential recruitment of afferent populations rather than a single fixed response to EUS stimulation. We observed that increasing pulse duration substantially reduced both current and frequency thresholds required to induce urinary leakage. This finding is consistent with the classical strength–duration relationship, where increasing pulse duration increases charge per pulse and enhances the probability of recruiting afferent fibers. Because pudendal afferents embedded within and around the EUS mediate urethra–bladder reflexes, increased charge delivery likely expands the population of activated sensory fibers. Notably, pulse duration shifted the entire current–frequency threshold surface rather than acting independently, demonstrating strong interdependence among stimulation parameters.

The functional consequences of urethral afferent activation are known to be state dependent. Danziger and Grill showed that urethral sensory feedback evokes different lower urinary tract reflexes depending on bladder volume and neural state [21]. At low bladder volumes, urethral flow preferentially evokes a guarding reflex characterized by EUS activation, whereas at higher volumes the same stimulus triggers an augmenting reflex that facilitates bladder contraction. Their work identified a distinct bladder volume threshold at which reflex behavior switches between continence and micturition modes. Although our experiments were conducted under isoflurane anesthesia, the parameter-dependent leakage observed here is consistent with recruitment of excitatory urethrovesical reflex pathways once sufficient afferent activation is achieved. Robain et al. demonstrated in awake ewes that urethral flow alone can reliably evoke robust detrusor contractions when bladder volume exceeds a threshold [20]. These responses were abolished by urethral anesthesia, confirming their sensory origin. Interestingly, even relatively small urethral flow rates were sufficient to trigger bladder contractions in the awake state. In our preparation, stimulation-induced leakage may allow small amounts of fluid to enter the urethra, potentially reinforcing bladder contraction via this urethrovesical reflex mechanism. While our study does not directly isolate this contribution, the observed leakage responses are physiologically consistent with previously described flow-evoked reflex activation.

With respect to aging, Oshiro et al. examined age-related changes in urethral smooth muscle and EUS function in rats [22]. They reported reduced urethral contractile responses and impaired coordination between detrusor contraction and sphincter relaxation in older animals, resulting in decreased voiding efficiency. These changes reflect alterations in urethral smooth muscle responsiveness and neuromuscular function rather than a fundamental reorganization of reflex circuitry. Consistent with those findings, we observe modest shifts in absolute leakage thresholds across age groups, yet the fundamental relationships among current, frequency, and pulse duration remain preserved. This suggests that while aging may alter baseline urethral mechanics and excitability, the core parameter-response dynamics governing afferent recruitment remain stable.

Several limitations should be acknowledged. All experiments were performed under isoflurane anesthesia, which can modulate reflex responsiveness and may elevate activation thresholds compared to awake conditions [21]. Additionally, urine leakage was used as a functional endpoint and does not differentiate between increased detrusor pressure and reduced outlet resistance. Future studies incorporating cystometric pressure recordings will help clarify the relative contributions of bladder activation and sphincter modulation. Finally, expanded investigation in disease models will be necessary to determine therapeutic applicability.

Overall, these findings provide mechanistic insight into how EUS stimulation parameters govern reflex bladder responses. By demonstrating the interdependent effects of pulse duration, current amplitude, and frequency, this study establishes a quantitative framework for parameter optimization in EUS-targeted neuromodulation. Such parameter-specific understanding is essential for developing precision stimulation strategies aimed at restoring bladder–sphincter coordination in individuals with neurogenic lower urinary tract dysfunction.

## V. CONCLUSION

This study demonstrates that pulse duration, current amplitude, and stimulation frequency interact synergistically to determine urine leakage thresholds during EUS stimulation. Rather than acting independently, these parameters exhibit strong interdependence, with changes in pulse duration shifting the current–frequency combinations required to elicit leakage. Animal age and sex exhibit only modest shifts in absolute thresholds and do not alter the fundamental parameter–response relationships. From a translational standpoint, these findings provide a clear framework for developing neuromodulation protocols that achieve desired bladder outcomes with minimal total charge delivery. In particular, optimizing stimulation parameters may reduce tissue fatigue, limit stimulation induced damage, and improve energy efficiency in future implantable devices designed to restore lower urinary tract function. Future work will include testing in awake, behaving animals with integrated cystometric pressure measurements to evaluate long-term stimulation effects, along with studies in disease models such as DSD to assess clinical therapeutic potential. Collectively, our findings support the development of next-generation, precision-controlled neuromodulation platforms for restoring bladder–sphincter coordination in NLUTD.

