## Supplemental File for "Quantitative Analysis of External Urethral Sphincter Stimulation Parameters for Modulating Urinary Output"

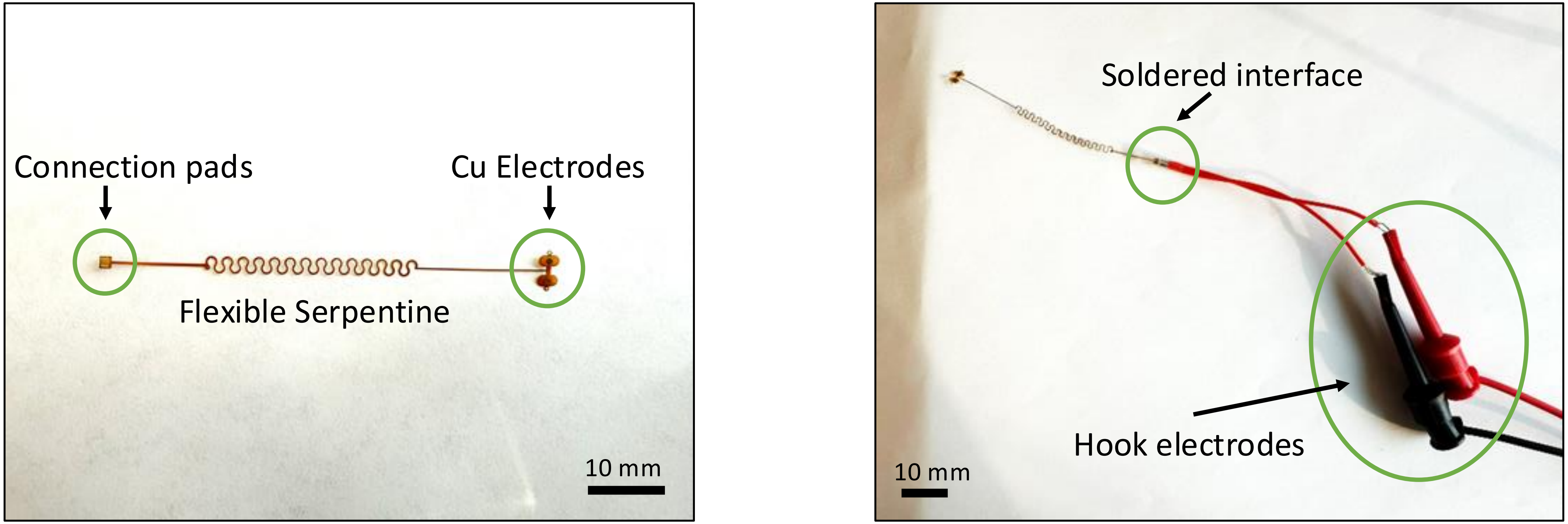


(a) (b)

**Supplementary Figure 1. Serpentine Connection with Electrode (A)** Copper electrode with flexible serpentine connection; **(B)** Electrode connection pads connected with copper wire by soldering, and another side connected with hook electrodes

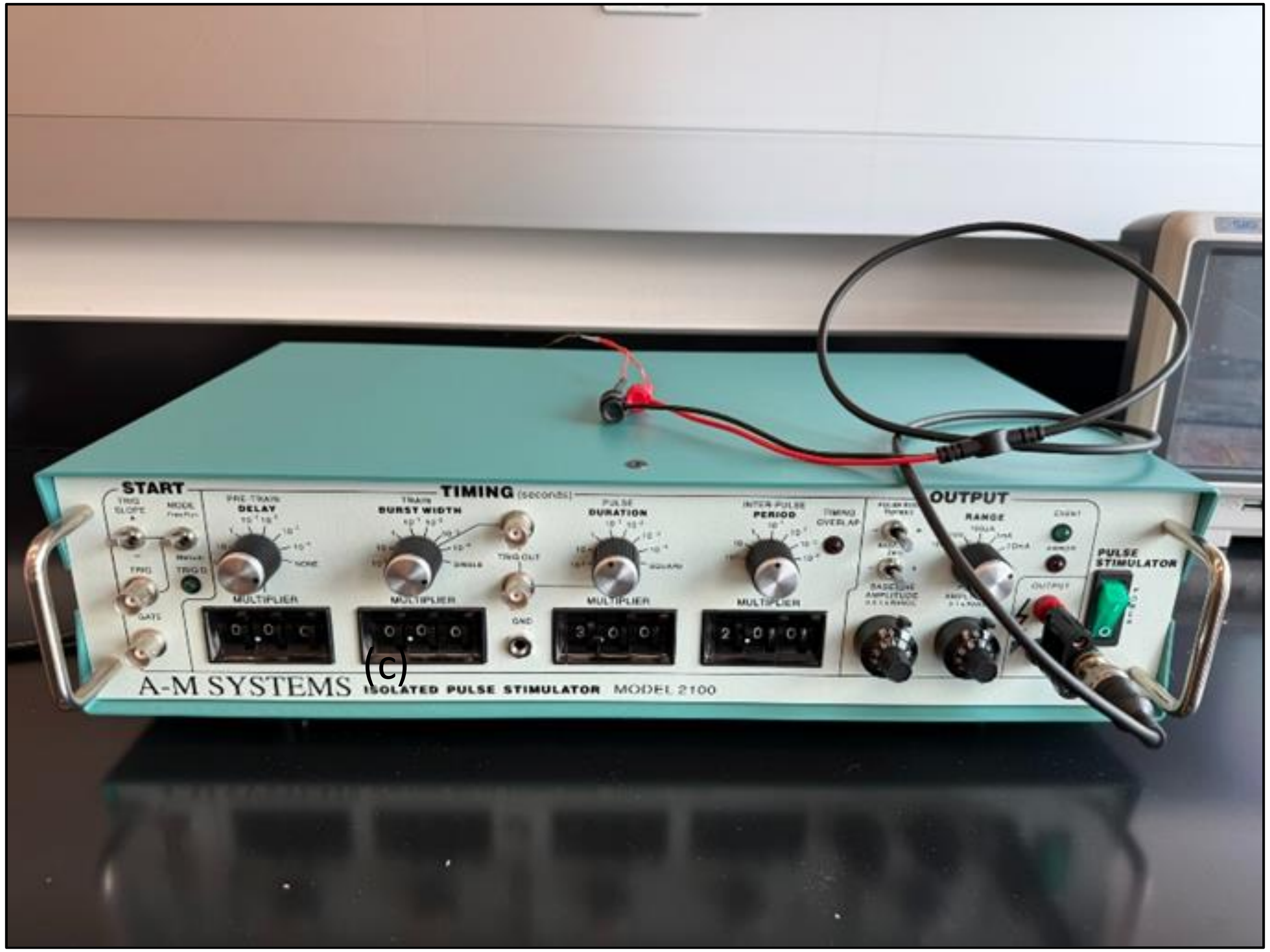


**Supplementary Figure 2.** Electrical Biphasic Pulse Stimulator

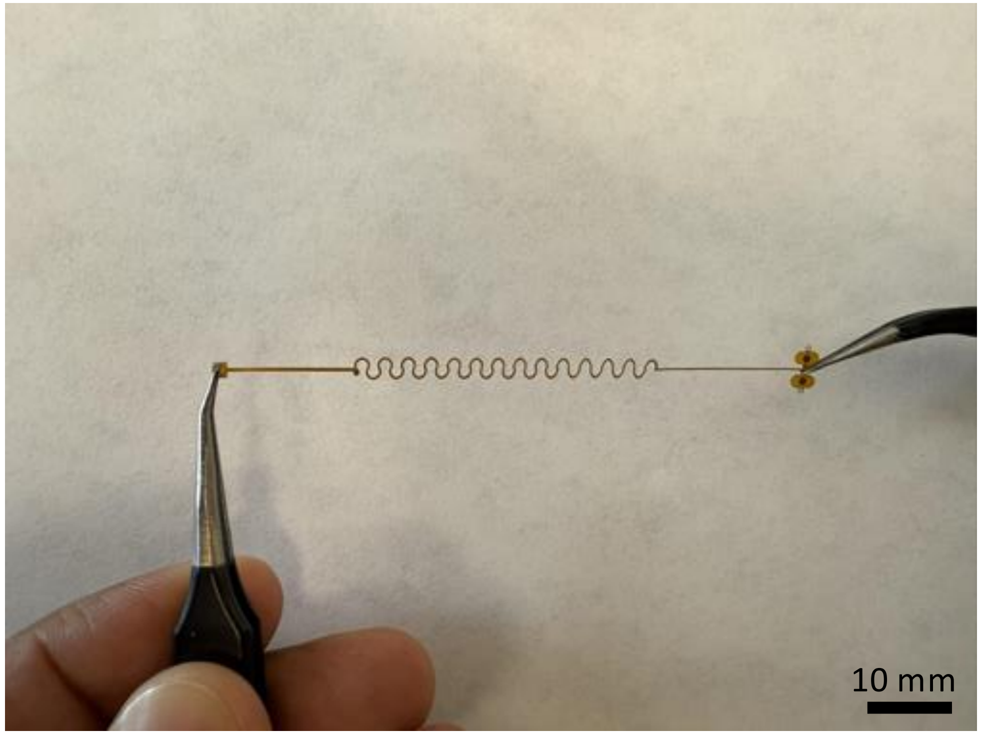


(a) Before stretching

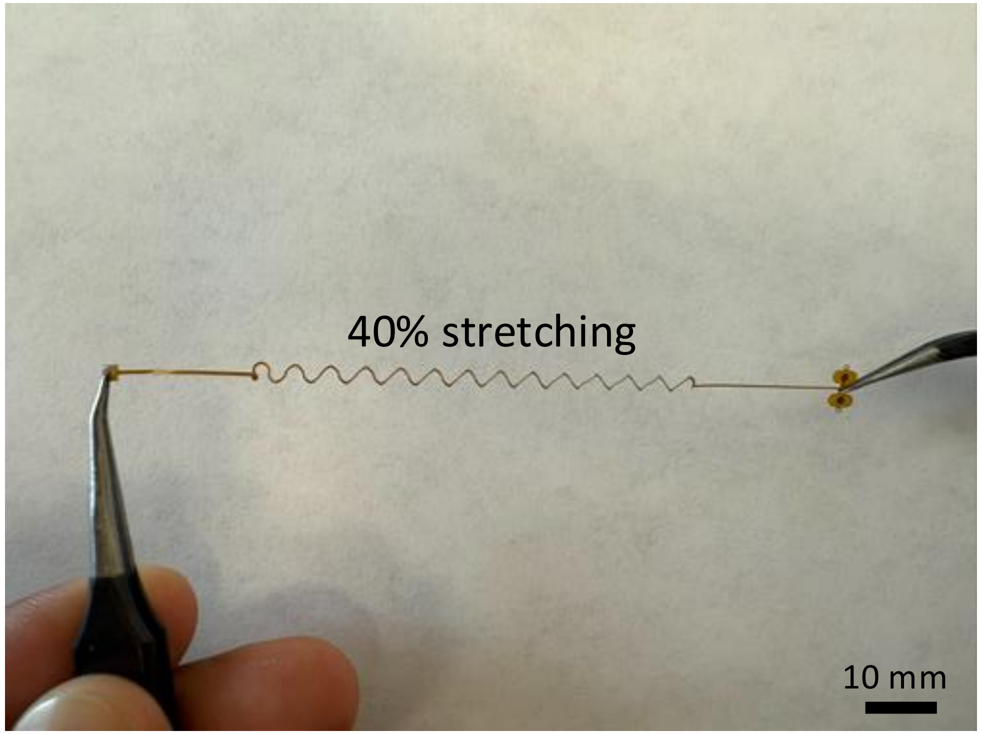


(b) After stretching

**Supplementary Figure 3. Electrode Serpentine Flexibility (A)** Before stretching; **(B)** After stretching.
